# Branch angle responses to photosynthesis are partially dependent on *TILLER ANGLE CONTROL 1*

**DOI:** 10.1101/251017

**Authors:** Jessica M. Guseman, Christopher Dardick

## Abstract

Light serves as an important environmental cue in regulating plant architecture. Previous work had demonstrated that both photoreceptor-mediated signaling and photosynthesis play a role in determining the orientation of plant organs. *TILLER ANGLE CONTROL 1* (*TAC1*) was recently shown to function in setting the orientation of lateral branches in diverse plant species, but the degree to which it plays a role in light-mediated phenotypes is unknown. Here, we demonstrated that *TAC1* expression was light dependent, as expression was lost under dark or far-red growth conditions, but did not display any clear diurnal rhythm. Loss of *TAC1* in the dark was gradual, and experiments with photoreceptor mutants indicated this was not dependent upon Red/Far-Red or Blue light signaling, but partially required the signaling integrator *CONSTITUTIVE PHOTOMORPHGENESIS 1* (*COP1*). Over-expression of *TAC1* partially prevented the narrowing of branch angles in the dark or under Far-Red light. Treatment with the carotenoid biosynthesis inhibitor Norflurazon or the PSII inhibitor DCMU led to loss of *TAC1* expression similar to dark or far-red conditions, but surprisingly expression increased in response to the PSI inhibitor Paraquat. Our results indicate that *TAC1* plays an important role in modulating plant architecture in response to photosynthetic signals.

**HIGHLIGHT:** Branch angles narrow in darkness or under far-red light. This response is partially mediated by *TAC1* which responds to photosynthetic signals, providing a key link between photosynthesis and plant architecture.

## INTRODUCTION

Plant architecture is intimately connected to light. It both influences the ability of the plant to intercept light and adjusts in response to light conditions. Architectural parameters such as organ angles, organ numbers, and branch lengths influence the quantity of light a plant can capture. For example, increased leaf number increases photosynthetic surface area, larger plant size and longer branches can allow plants to avoid shade from their neighbors, and leaf angle changes with respect to the angle of sunlight influence the amount of light captured (Osada and Hiura, 2017). In turn, changes in light quality and quantity result in the modification of these parameters. Growing plants under shaded conditions, for example, results in phenotypes characteristic of shade avoidance syndrome, including upward leaf movement, accelerated elongation of plant organs, and fewer shoot branches (Casal, 2012). In addition to these, shade also leads to more vertically oriented branches in Arabidopsis (Roychoudhry et al., 2017).

Lateral organ orientation, or angle, is an important aspect of plant architecture that has been connected to multiple light signaling pathways. Recent work addressing neighbor detection demonstrated that petiole angle altered in response to FR light detection at the leaf margin (Pantazopoulou et al., 2017). These studies showed a connection between R/FR light signaling and architecture. Early work defining gravitropic set point angle, the angle at which organs grow with respect to gravity, identified a regulatory role for photosynthesis using *Tradescantia* as a model (Digby and Firn, 2002). However, beyond this study little work has been done to elucidate the connection between photosynthesis and branch angles.

Studies to determine the endogenous genetic components underlying lateral organ orientation identified loci associated with narrowed angles in Rice, Maize, and Brassica (Yu et al., 2007; Ku et al., 2011; Li et al., 2017). A gene repeatedly identified in these studies, *TILLER ANGLE CONTROL 1* (*TAC1*), has been shown to regulate lateral branch angle in Arabidopsis, peach, and plum (Dardick et al., 2013; Hollender et al., in press). Loss of *TAC1* expression, through mutation or silencing, results in more vertical organ orientation in tillers, branches, leaves, and pedicels. In peach canopies this led to increased rate of carbon accumulation, as the changes in canopy shape allowed increased light penetrance (Glenn et al., 2015). *TAC1* belongs to the IGT family, named for a shared amino acid motif, which also contain *LAZY* and *DEEPER ROOTING* (*DRO*) genes (Hollender and Dardick, 2015). Members of the *LAZY* and *DRO* clades have recently been reported to influence both shoot and root organ orientation via changes in gravity response upstream of auxin transport (Yoshihara et al., 2013; Ge and Chen, 2016; Guseman et al., 2016; Taniguchi et al., 2017; Yoshihara and Spalding, 2017). Currently, little is known about the regulation of IGT genes, however *LAZY1* expression in maize was reported to be lower under light conditions (Dong et al., 2013).

Here we address the hypothesis that *TAC1* is involved in light regulation of lateral branch angles. Our results show that *TAC1* exhibits light dependent gene expression, which correlates with narrowed branch angles in response to prolonged growth in darkness. Constitutive expression of *TAC1* could partially, but not fully rescue changes in lateral branch orientation. *TAC1* expression was not dependent upon known photoreceptor signaling pathways, but partially required a fully functional *CONSTITUTIVE PHOTOMORPHGENESIS 1* (*COP1*) gene. Using various photosynthetic inhibitors, we found that *TAC1* expression was abolished when treated with Norflurazon (NF) and 3-(3,4-Dichlorophenyl)-1,1-dimethylurea (DCMU) and increased in response to Paraquat (PQ) treatment, suggesting that *TAC1* is a target of photosynthetic signals to alter the angle of organs in response to persistent changes in light exposure.

## MATERIALS AND METHODS

### Plant material and growth conditions

The Columbia (Col-0) and Landsberg erecta (Ler) ecotypes were used as WT lines in all experiments. Signaling mutants *phyAB* and *phyABDE* (Hu et al., 2013), *cry1;cry2* (Mockler et al., 1999), *phot1;phot2* (Kinoshita et al., 2001), *cop1-6* (Ang and Deng, 1994), *pifQ* (Lilley et al., 2012) and *hy5;hfr1;laf1* (Jang et al., 2013) were previously described. For phenotyping and expression studies, seeds were surface sterilized and sown on square plates containing half-strength MS and 0.8% bactoagar and grown vertically. Once sown, seedlings were stratified at 4°C in the dark for 2 days, then placed in growth chambers at 20°C with a 16-h light/8-h dark photoperiod (~100 µmol m^2^ sec^-1^).

### Branch angle measurements

For shoot branch angles, seedlings were grown for 2 weeks on plates, then transplanted into 4-inch pots containing Metromix 360 soil (Sun-Gro Horticulture, http://www.sungro.com) and grown until bolting (~6–7 inches in height). Plants were then transferred to continuous light or dark conditions for 72 hours. Bolts were then photographed and pressed. Images were taken using a Canon EOS Rebel T3 camera (http://global.canon/en/index.html). Angles were manually calculated by measuring the angle of the tangent of each lateral branch point, with respect to the upper main stem.

### RNA extraction and quantitative real time PCR

Arabidopsis seedlings were grown on vertical plates for 10-14 days. Four biological replicates were used. Each biological replicate consisted of a plate of 10-12 seedlings. Arabidopsis RNA was extracted using a Directzol RNA Extraction Kit (Zymo Research, http://www.zymoresearch.com). qPCR was performed as previously described by Dardick et al. (2010). Briefly, each reaction was run in triplicate using 50 ng of RNA in a 12µl reaction volume, using the Superscript III Platinum SYBR Green qRT-PCR Kit (Invitrogen, now ThermoFisher Scientific, https://www.thermofisher.com). The reactions were performed using a 7900 DNA sequence detector (Applied Biosystems, now ThermoFisher Scientific, https://www.thermofisher.com). Quantification for Arabidopsis samples was performed using a relative curve derived from a serially diluted standard RNA run in parallel. *UBC21* was used as an internal control to normalize expression in light experiments, and *IPP2* was used for circadian experiments.

### Light and time-course experiments

For light experiments, plants were grown for 10 days on vertical plates in 16:8 long day light conditions in a growth chamber before transfer to experimental light conditions. For comparisons between light and dark, plates were moved to chambers with either continuous light or continuous dark conditions for 72 hours, then whole seedlings were collected and flash frozen at 10am (ZT4). For comparisons between light colors, plates were moved to chambers with continuous white, red, blue, or far red light for 72 hours and whole seedlings were collected at 10am (ZT4). Matching growth chambers fitted with white, red, blue, and far-red LED lamps from PARsource (http://parsource.com) were used for light color experiments. For circadian experiments, seedlings were grown for 10 days in 12L:12D light cycles, then transferred to continuous light and collected every 4 hours for 84 hours. For adult phenotypes, plants were grown on soil for 5-6 weeks, until bolts reached 4-6 inches in height. Then plants were transferred to continuous W or FR light conditions for 72 hours then imaged and collected.

### Chemical treatments

For sucrose experiments, plants were germinated and grown on 0.5x MS plates for 10 days, then transplanted to plates containing 1% sucrose. Plates were then moved to continuous light or dark conditions for 72 hours and collected at 10am (ZT4). For photosynthesis experiments, plants were grown on vertical MS plates for 7 days, then transplanted onto media containing either Norflurazon (5uM), DCMU (10uM), Paraquat (1uM), or mock (water). Plates were then moved to continuous light or dark conditions for 5 days and collected at 10am (ZT4). For treatment of adult plants, Arabidopsis were grown for 5-6 weeks until bolts reached 4-6 inches in height.

### Chlorophyll Fluorescence Imaging

All chlorophyll fluorescence was measured using the Maxi-Imaging-PAM Chlorophyll Fluorometer (Walz, Effeltrich, Germany). Maximum PSII quantum yield (Fv’/Fm’) was determined using an actinic light pulse (1500 μmol m-2 s-1). Average Fv’/Fm’ values were calculated for a similar area of interest for 6 seedlings on each chemical treatment, using the Maxi-Imaging-PAM software.

## RESULTS

### *TAC1* expression is lost under extended continuous dark conditions

To address whether *TAC1* plays a role in light regulation of organ angle, we initially screened the promoter region upstream of *TAC1* for the occurrence of light-related cis-elements (Fig 1A). Using a cis-element database (AGRIS AtcisDB, http://arabidopsis.med.ohio-state.edu/AtcisDB/), we identified several elements, including GATA motifs, a G-box, T-boxes, and AtMYC2 binding sites. Next, we tested the response of *TAC1* expression to plant growth in continuous dark for 72 hours. *TAC1* expression was lost while that of a control gene, *UBC21*, was unaffected (Fig 1B). To determine the dynamics of this expression loss, we performed a time course experiment over a 72 hour period of continuous dark. Expression levels gradually declined over time, reaching their lowest values by 48 hours (Fig 1C). Plants grown for 72 hours in continuous dark and then returned to continuous light showed similar expression dynamics. Expression began to increase around 4 hours once transferred back into the light, but did not return to normal levels until 48 hours (Fig 1D). To address whether *TAC1* exhibits a diurnal rhythm, we performed a circadian time course, transferring plants previously entrained to a 12L:12D light cycle to continuous light conditions. *TAC1* expression did not exhibit a clear rhythm (Fig 1E). Taken together, the data suggest *TAC1* expression is dependent on light, but with gradual response dynamics.

**Figure 1.**
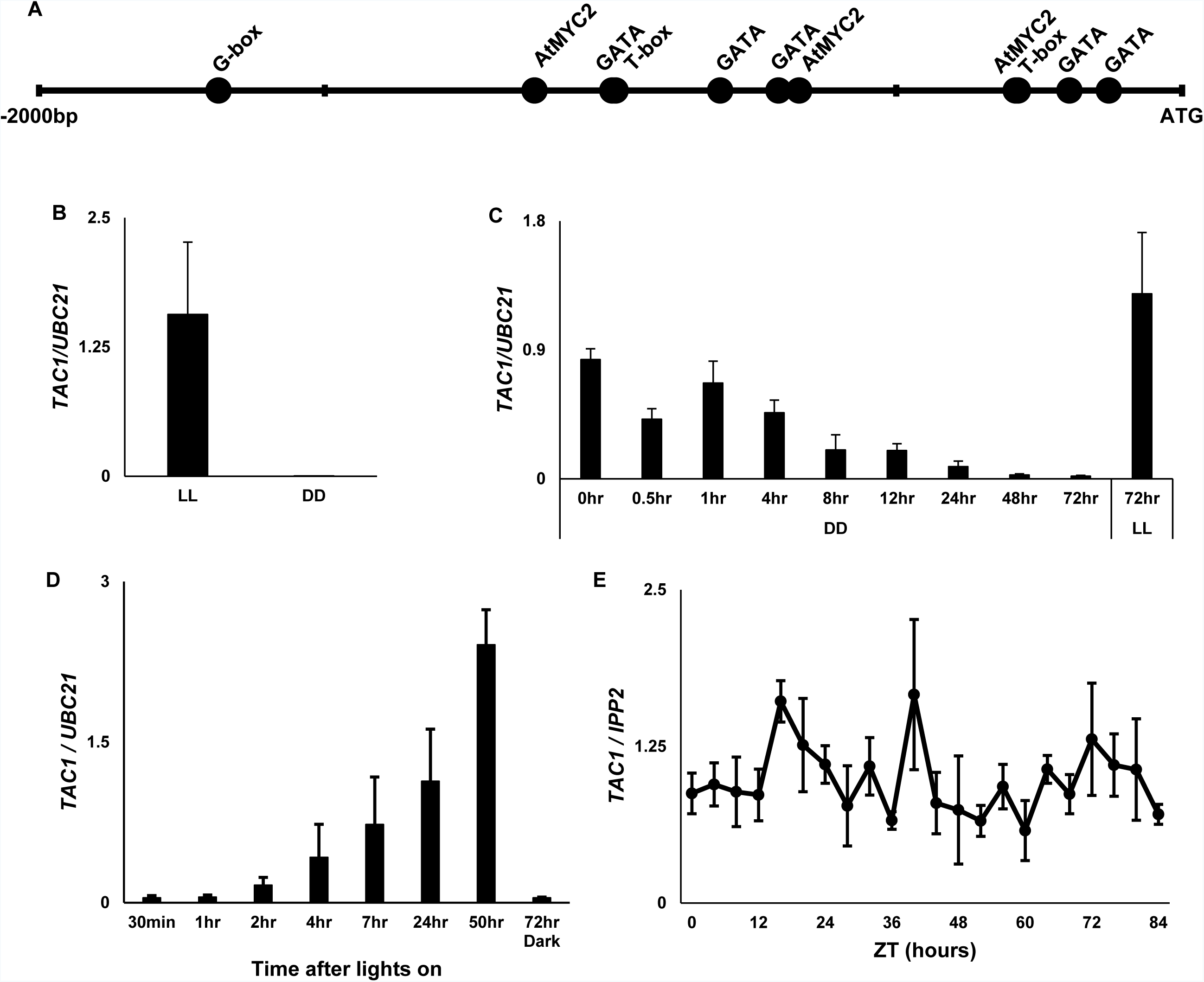
*TAC1* expression is light dependent. A. Promoter analysis of *TAC1* reveals light-related motifs. B. Quantitative-RT-PCR shows dramatic reduction in *TAC1* expression in wild-type seedlings grown in continuous dark for 3 days, as compared to continuous light. C. Time course qRT-PCR data from plants moved to continuous dark show that complete loss of *TAC1* expression occurs between 24-48 hours in dark. D. Time course qRT-PCR data taken from plants moved from 3 days continuous dark to continuous light demonstrate that TAC1 expression returns to original levels after 24-48 hours in light. E. Plants transferred to continuous light maintain *TAC1* expression and do not show a clear circadian rhythm. Error bars represent SD.

### Lateral branch angles narrow in response to growth in 72h of continuous dark

To test whether the loss of *TAC1* expression in dark conditions correlated with changes in Arabidopsis branch angle phenotypes, we grew adult plants in continuous dark for 72 hours. Lateral branch angles of wild-type plants significantly narrowed by about 10 degrees compared to continuous light-grown controls (Fig 2). Plants overexpressing *TAC1* (*35S::TAC1*) still showed narrowed branch angles in dark conditions but not to the same degree as Col, suggesting there are *TAC1*-dependent and *TAC1*-independent pathways influencing this process. *tac1* mutants plants exhibited narrow angles, similar to dark-grown wild-type plants, in both light and dark conditions.

**Figure 2.**
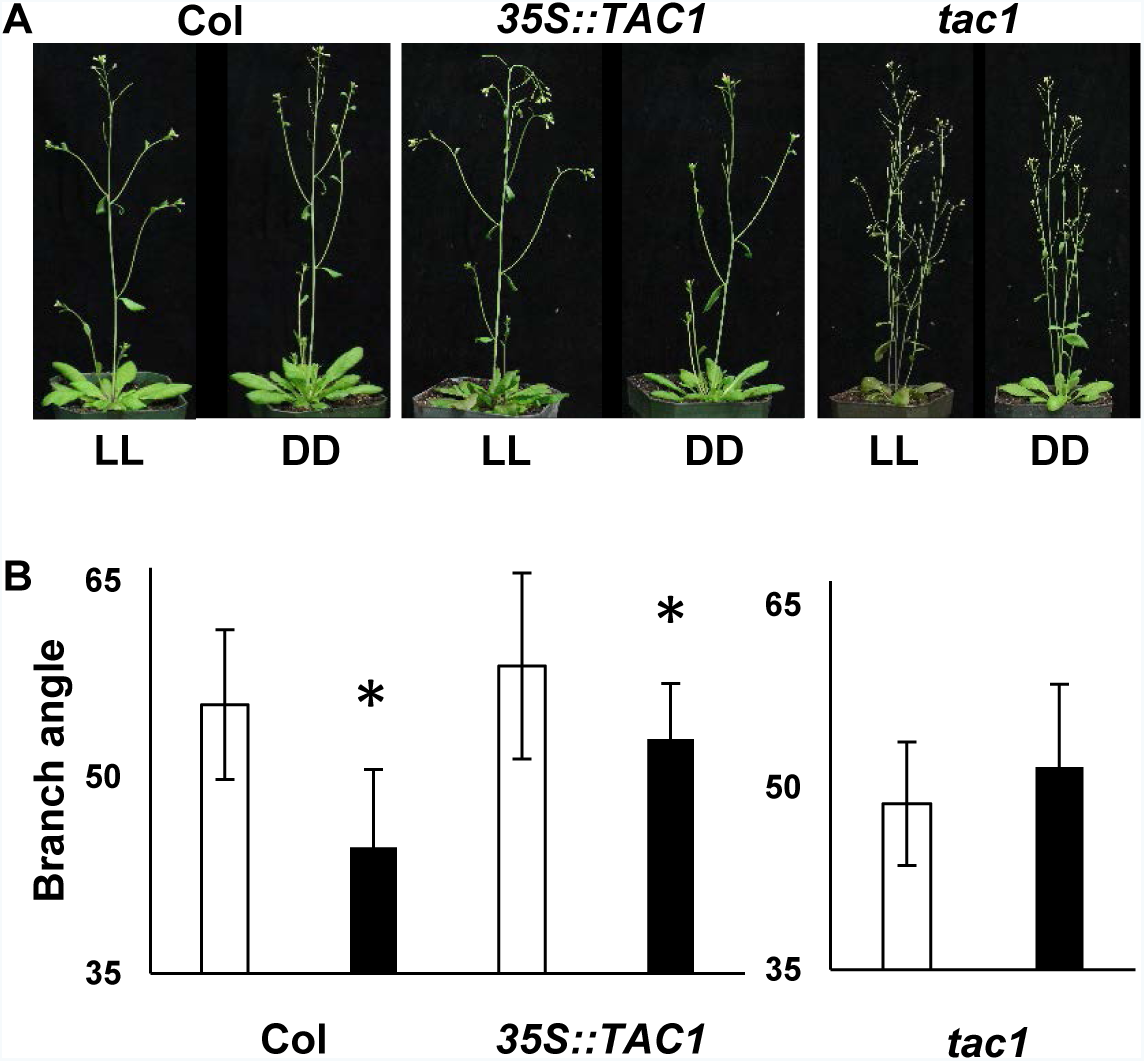
Dark-grown Arabidopsis plants exhibit vertically oriented branch growth. A. Wild-type (Col), *35S::TAC1*, and tac1 plants grown in continuous light or dark for 72h. B. Quantification shows a significant decrease in wild-type and *35S::TAC1* dark-grown lateral branch angle with respect to the upper stem. Error bars represent SD.

### *TAC1* is lost in FR light, does not require *phys*, *crys*, or *phots*, but is reduced in a weak *cop1* mutant background

We next sought to determine which aspects of light were required for *TAC1* expression. First, we tested the requirement for specific light wavelengths, growing plants in 72 hours of continuous white (W), red (R), blue (B), or far-red (FR) light (Fig 3B). In comparison to growth in W light, *TAC1* expression was not significantly different under R, and elevated slightly, about two-fold, under B light. Under FR light, the response was similar to growth in darkness, with very low levels of expression. We tested whether FR treatment reduced expression in adult plants and led to similar changes in branch angles observed in dark-grown plants. Adult wild-type and *35S::TAC1* plants were grown in continuous W and FR light for 72 hours, then plants were imaged, lateral apices were collected, and angles at branch points were measured. Compared to W light, FR-grown plants showed loss of *TAC1* expression, similar to dark conditions, and branch angles narrowed by about 8 degrees (Fig 3 C-D). Contrary to this, plants containing a *35S::TAC1* construct did not show a reduction in *TAC1* expression in FR light. Branch angles narrowed slightly but not to the same degree as Col plants in response to FR light. These results were consistent with dark experiments and confirms there are likely both *TAC1*-dependent and independent mechanisms for regulating branch angles in response to changes in light.

**Figure 3.**
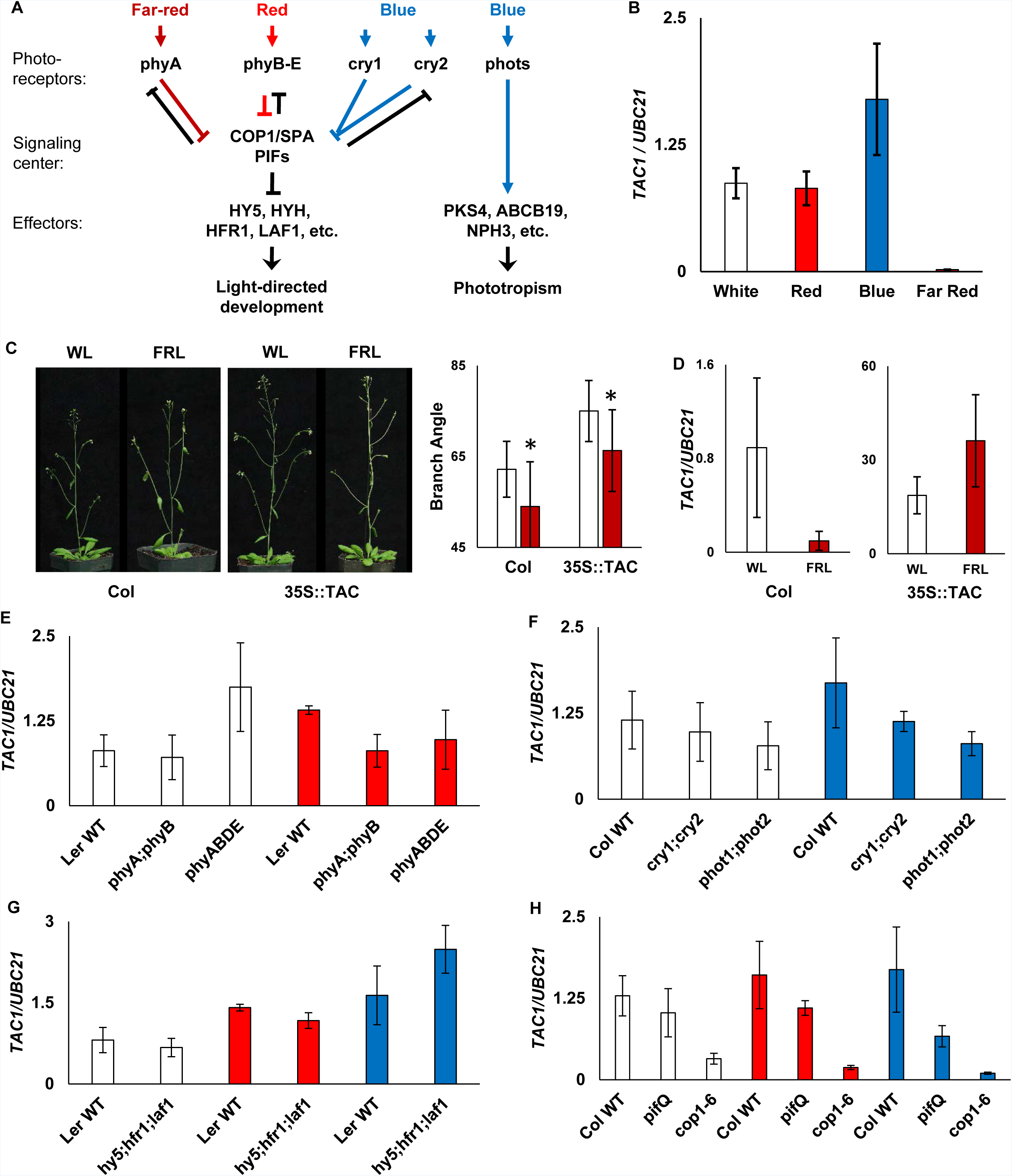
*TAC1* expression is decreased in FR light and *cop1* mutant background. A. Model of phytochrome, cryptochrome and phototropin light signaling pathways. Adapted from Lau and Deng, 2012. B. qRT-PCR expression data in W, R, B, FR light shows *TAC1* is downregulated in FR conditions. C. Representative WT and *35S::TAC1* plants grown in W and FR light for 3 days, and quantified branch angles. n=8 plants per treatment D. *TAC1* expression in Col and *35S::TAC1* branch apices after 3 days of W or FR light treatment. E. *TAC1* expression in Col WT, cryptochrome, and phototropin mutants, grown in continuous white or blue light for 3 days. F. *TAC1* expression in Ler WT and phytochrome mutants, grown in continuous white or red light for 3 days. G-H. *TAC1* expression in WT and mutants involved in both red and blue light signaling pathways, *cop1*, *pifQ*, and *hy5;hfr1;laf1*, grown in continuous white, red, or blue light for 3 days. Error bars represent SD.

The findings prompted us to explore two potential mechanisms by which *TAC1* expression could be regulated by light: first, that *TAC1* expression requires either R or B light via photoreceptor signaling, or second, that *TAC1* expression is controlled by another light-related process such as photosynthesis. To test the first, we looked at *TAC1* levels in different photoreceptor and light-signaling mutant backgrounds, grown under W, R, or B light. While there were small, but significant changes in expression in some photoreceptor mutant backgrounds (Figs 3E and F), none of these changes could explain the loss of *TAC1* observed in the dark. For example, if phytochromes were required for *TAC1* expression, then loss of *TAC1* would be expected in a *phy* mutant background grown under R light. There was a relatively small decrease in *TAC1* expression in the *phyAB* mutant in R light, however this does not mimic dark-growth results, and the quadruple *phyABDE* mutant did not show a similar effect (Fig 3E). Similarly, there was a small but significant loss of *TAC1* expression in the *phot1;phot2* background as compared to Col WT in B, however not enough to explain loss of gene expression in the dark (Fig 3F). In addition, we used several mutants downstream of both R/FR and B light signaling pathways: a weak *cop1* allele, a triple *hy5;hfr1;laf1* mutant and the *pif1;pif3;pif4;pif5* (*pifQ*) mutant (Figs 3G and H). Similar to the photoreceptor mutants, we saw relatively minor or insignificant changes in *TAC1* levels in *pifQ* and *hy5;hrf1;laf1* mutant backgrounds. To the contrary, we saw a larger and significant reduction in expression in *cop1-6* mutants. Together, the data suggest that different aspects of R/FR and B light signaling may influence *TAC1* expression to a small degree, but do not explain the loss of expression in dark-grown plants.

### Exogenous sucrose does not rescue loss of *TAC1* in the dark

Sucrose has been reported to have an effect on lateral organ angle (Willemoes et al., 1988), and dark-grown plants have decreased photosynthetic efficiency, and thus produce less photosynthate. To test whether *TAC1* expression is dependent on the products of photosynthesis, we grew plants on media supplemented with sucrose and exposed these to continuous light and dark conditions (Fig 4A). Gene expression was similar when supplemented with sucrose in both conditions, demonstrating that exogenous sucrose was not sufficient to attenuate the loss of *TAC1* expression in the dark. This suggests that sucrose-mediated alteration of organ angle is *TAC1*-independent.

**Figure 4.**
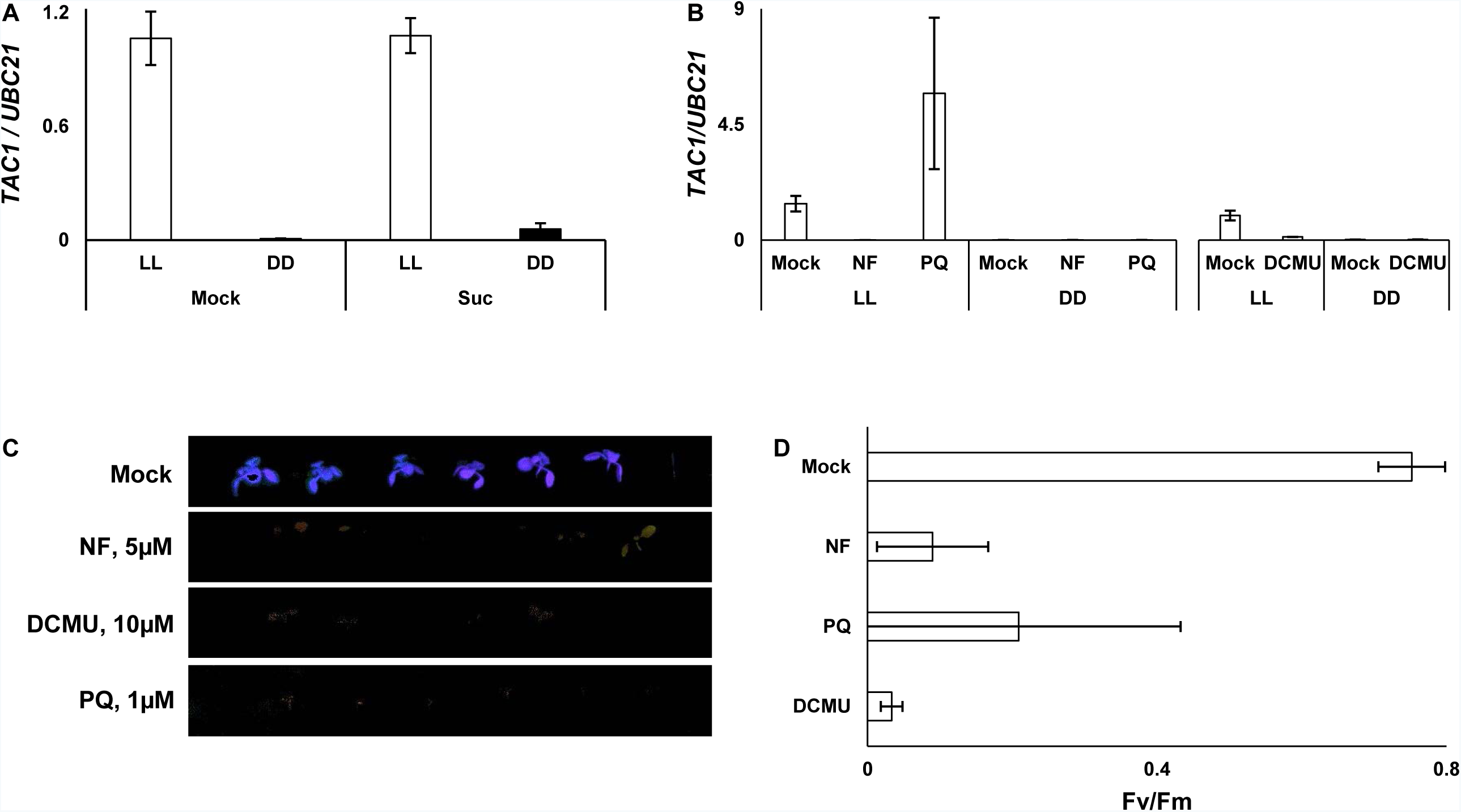
Sucrose and photosynthesis inhibitors have differential effects on *TAC1*
expression. A. *TAC1* expression in plants grown on media with and without sucrose show no significant different between treatments. B. *TAC1* expression in plants grown in 72h continuous light or dark after transplant to media containing NF, LM, PQ, or a mock control. Expression is decreased when treated with NF, and increased when treated with LM or PQ. C. Cholorphyll fluorescence imagine of plants treated with NF, LM, and PQ. D. Quantified photosynthetic efficiency, measured as average Fv/Fm, in plants treated with NF, LM, and PQ.

### Photosynthetic inhibitors have differential effects on *TAC1* expression

To test if *TAC1* expression is regulated by photosynthetic activity, we treated plants with a series of photosynthesis inhibitors. Each of these inhibitory chemicals impairs photosynthesis through different pathways. Treatment with norflurazon (NF) inhibits carotenoid biosynthesis, allowing for the formation of triplet chlorophyll and subsequent photooxidating damage within the chloroplast (Gray et al., 2003). DCMU specifically inhibits electron transport by blocking the plastoquinone binding site of Photosystem II. In contrast, Paraquat (PQ), also known as methyl viologen, acts by shunting electrons from Photosystem I, and producing high levels of reactive oxygen species (ROS). 7 day-old seedlings transferred to media supplemented with these photosynthetic inhibitors were grown in continuous light or dark and measured for photosynthetic efficiency (Fv/Fm) and *TAC1* gene expression (Fig 4B-D). Treatment with NF led to decreased photosynthetic efficiency, as measured by chlorophyll fluorescence imaging, and abolished *TAC1* expression in the light, mimicking the effect observed in dark-grown plants (Fig 4B-D). DCMU treatment resulted in near total loss of chlorophyll fluorescence, and treated plants showed a similar decrease in *TAC1* expression as with NF treatment. PQ treatment displayed an inconsistent reduction in PSII efficiency, but led to variable but significant increases in *TAC1* expression (Fig 4B-D). All plants grown under continuous dark conditions exhibited loss of *TAC1*, regardless of treatment (Fig 4B).

## DISCUSSION

Lateral organ angle is strongly tied to light capture, which has important implications for plant productivity and competition. Previously, a connection between photosynthesis and branch angle was described in *Tradescantia* by Digby and Firn (2002). We provide evidence that *TAC1* is a target of photosynthetic signals, and is partially required for the changes in lateral branch angles that are driven by photosynthesis. Arabidopsis grown in continuous darkness exhibited more vertically oriented lateral branches, phenocopying a *tac1* mutant phenotype. *TAC1* expression in dark-grown plants was abolished after 24-48h, suggesting that this mechanism is in place to induce vertical growth when branches are subjected to extended periods of darkness. Consistent with this, *Tradescantia* plants treated with the photosynthetic inhibitor DCMU grew upward, mimicking their growth in dark conditions (Digby and Firn, 2002). Treatment with NF results in triplet chlorophyll formation, and also decreases nuclear gene expression involved in multiple photosynthetic processes, including the light harvest complex, electron transfer chain, photosystem II oxygen-evolving complex, and the reductive pentose phosphate pathway, (Gray et al., 2003), effectively reducing function of multiple early steps in photosynthesis. The loss of *TAC1* expression in response to NF treatment may suggest that photosystem II function is required. PQ effectively reduces photosystem I function, later in photosynthesis, and also generates ROS production. The increase of expression in response to PQ suggests that *TAC1* does not require photosystem I, and may be sensitive to ROS signaling. Taken together, it is likely that *TAC1* functions downstream of photosynthesis as a regulator of branch angle.

Both sucrose treatment and photoreceptor-mediated light signaling play roles in setting lateral organ angles (Willemoes et al., 1988; Pantazopoulou et al., 2017; Roychoudhry et al., 2017). However, neither had a strong influence on *TAC1* expression. Growth in FR light both decreased *TAC1* expression and led to narrowed branch angles, but *TAC1* remained relatively unaffected by R/FR signaling components. Recent work demonstrated that *PIF4* is not required for shade-induced reduction in lateral branch angle (Roychoudhry et al., 2017). Our finding that *TAC1* expression is unchanged in a *pifQ* mutant background is consistent with this finding. Blue light led to elevated levels of TAC1 in several experiments. However, large increases in expression, in the case of 35S::TAC1 plants, had little effect on increasing branch angle. Together, these data suggests that the influence of both sucrose, and B and R/FR light signaling on organ orientation is largely *TAC1*-independent.

Of the light signaling mutants tested, *cop1-6* mutants had the strongest effect on *TAC1* gene expression. However, the effect of *COP1* appears to be independent of phytochrome or cyptochrome-mediated signaling, as other mutants within these pathways exhibited little to no change. Recent work has implicated *COP1* in chloroplast retrograde signaling, revealing that *COP1* degrades *ABI4* in the light during de-etiolation (Xu et al., 2016). The requirement of *COP1* coupled with the differential responses of *TAC1* expression to chemical inhibitors of photosynthesis raises the question whether *TAC1* is regulated by retrograde signaling. The data presented here suggests a possible signaling pathway from photosynthesis, through *COP1* and *TAC1* to regulate branch angles, and that *TAC1* may function as part of a feedback mechanism by which plants modify branch orientations to optimize light capture and photosynthetic efficiency.

## ACKNOWLEDGEMENTS

We would like to thank the labs of Kerry Franklin, Chentao Lin, Ken-ichiro Shimazaki, Xing Wang Deng, Jennifer Nemhauser, and In-Cheol Jang for providing seeds of the light signaling mutants. The work at AFRS was supported by Agriculture and Food Research Initiative Competitive grant 10891264 from the USDA National Institute of Food and Agriculture and by the National Science Foundation grant number 1339211.

## AUTHOR CONTRIBUTIONS

JMG designed experiments and performed analyses. JMG wrote the manuscript with help from CD.

